# A proposed workflow to analyze bacterial transcripts in RNAseq from blood extracellular vesicles of people with Multiple Sclerosis

**DOI:** 10.1101/2024.04.23.590754

**Authors:** Alex M. Ascensión, Miriam Gorostidi-Aicua, Ane Otaegui-Chivite, Ainhoa Alberro, Rocio del Carmen Bravo-Miana, Tamara Castillo-Trivino, Laura Moles, David Otaegui

## Abstract

**Motivation:** The taxonomical characterisation of bacterial species derived from genetic material blood, including reads derived from bacterial extracellular vesicles (bEVs) poses certain challenges, such as the proper discrimination of “true” reads from contaminating reads. This is a common issue in taxa profiling and can lead to the false discovery of species that are present in the sample. To avoid such biases a careful approximation to taxa profiling is necessary.

**Results:** In this work we propose a workflow to analyze the presence of bacterial transcripts as indicative of putative bEVs circulating in the blood of people with MS (pwMS). The workflow includes several reference mapping steps against the host genome and a consensus selection of genera based on different taxa profilers. The consensus selection is performed with a flagging system that removes species with low abundance or with high variation across profilers. Additionally, the inclusion of biological samples from known cultured species as well as the generation of artificial reads constitute two key aspects of this workflow.

**Availability:** The workflow is available at the following repository: https://github.com/NanoNeuro/EV_taxprofiling.

## Introduction

Extracellular vesicles (EVs) can be defined as membrane vesicles characterized by their small size, naturally produced as lipid bilayer vesicles that contain a cargo (DNA, RNA, protein, lipids, etc.). The term has been used to refer to eukaryotic EVs and they have been widely studied in human blood demonstrating their huge importance in health and disease [1]. However, eukaryotic vesicles are not the only ones that can be found in human biofluids. Bacterial EVs (bEVs) arise from the outer and inner membranes of gram-negative bacteria and cytoplasmic membranes of gram-positive bacteria through blebbing and lytic biogenesis pathways [2, 3]. Although bEVs coming from Gram-negative have been usually referred to as outer-membrane vesicles (OMVs), bEVs could involve all the vesicles produced by bacteria [4]. The presence of bEVs has been reported in human tissues such as saliva, feces and also in blood. Despite the potentially pathogenic component of these bEVs, they could also be a relevant communication tool between microbiota and the rest of our organism [5, 6]. The influence of microbiota in several diseases is widely reported, including neurological and autoimmune diseases. In the case of Multiple Sclerosis (MS), the dysbiosis of the microbiota has been clearly related to the disease, through several experiments in mice [7, 8] and through a huge collaborative effort that characterizes the microbiome in a large cohort [9]. In this context, this work aims to analyze the presence of bacterial transcripts as indicative of putative bEVs circulating in the blood of people with MS (pwMS).

## Material and methods

A summary of the methods is available in Figure 1.

**Fig. 1.**
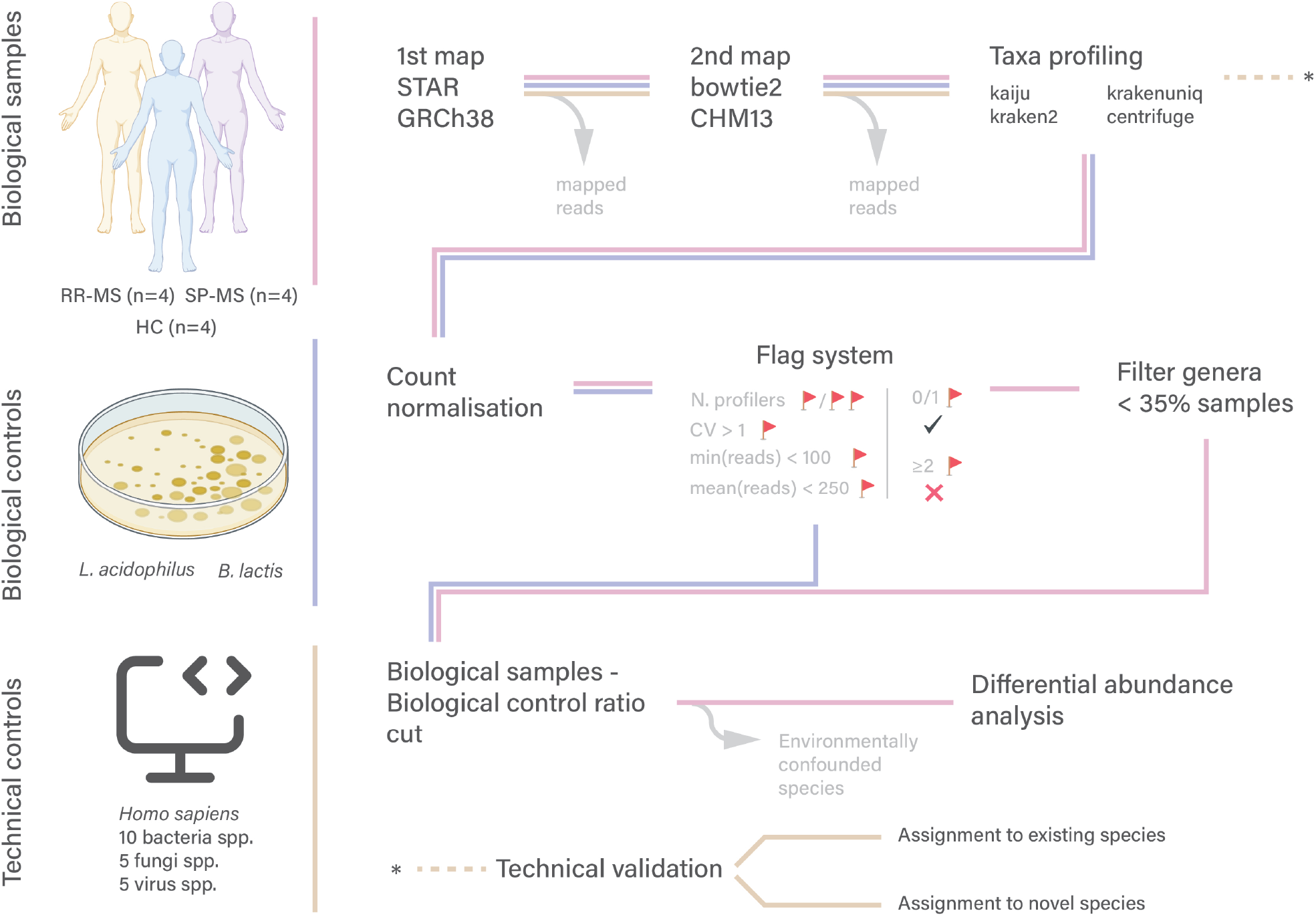
Summary of sample processing pipeline. Each of the steps is marked with differently colored lines depending on the sample they are referring to.

### Sample and EV extraction

As further described in Iparraguirre et al. [10], whole blood was obtained from MS patients (10 RR-MS and 10 SP-MS) and age-matched healthy controls (n = 8) at the Department of Neurology of the Donostia University Hospital (Supplementary Table S1). Peripheral blood was collected by venipuncture into EDTA tubes and centrifuged at 1258× g for 20 minutes to separate the plasma from the cellular fraction.

To isolate EVs, plasma aliquots were centrifuged at 13,000× g for 2 minutes at 4°C. The supernatant was transferred to a new tube and centrifuged again at 20,000× g for 20 minutes at 4°C to pellet EVs. The pellet was then resuspended with 100 µL DPBS (GIBCO, ThermoFisher, Waltham, MA, USA). RNA was isolated with Trizol LS (ThermoFisher, Waltham, MA, USA) as described in the work.

Finally, samples were pooled to achieve a minimum amount of 2 *µ*g RNA. rRNA was removed from the total RNA sample and cDNA libraries were prepared. Paired-end sequencing was performed using Illumina HiSeq X Ten, obtaining an average of 40–50 × 10^6^ reads per sample. Information about sample pooling is available at Supplementary Table S2.

### Read mapping to human reference

The reads from the samples were mapped to human references twice, based on the method described by Gihawi et al. [11]. A first mapping was performed with GRCh38 (Ensembl version 109) using the module *nf-core/rnaseq* v3.12.0 [12], with *STAR* v2.7.9a as mapper [13]. Unmapped reads were mapped again, this time with *bowtie2* v2.4.5 with the argument *–very-sensitive* and the CHM13 genome version (NCBI accession GCF 009914755.1).

### Taxa profiling of unmapped reads

Unmapped reads were then mapped with different profilers: *kaiju* v1.9.2 [14], *kraken2* v2.1.2 [15], *krakenuniq* v1.0.4 [16] and *centrifuge* v1.0.4 [17]. For each profiler, specific available databases were downloaded or created containing at least human, archaeal, bacterial, viral and fungal genomes. Specific settings for each profiler were as follows: for *kaiju*, e-value=0.0001 and minimum-length=41; for *kraken2*, confidence=0.90; for *krakenuniq*, hll-precision=16; and for *centrifuge*, minimum-hitlen=41. After the individual profiles were created, *taxpasta* v0.3.0 was used to standardize the outputs of the individual profilers into a common format [18]. The lines of the taxa were cut to genus level.

### Generation of control and **in silico** samples

Two types of additional samples were created to generate robust controls for comparison of sample results. On the one hand, control samples were generated by sequencing reads from EVs extracted from cultured *Lactobacillus acidophilus* and *Bifidobacterium lactis*. EVs were extracted by performing an initial centrifugation at 13,000*×*g for 10 minutes at 4°C; then were filtered via a 0.22 *µ*m filter and recentrifuged at 200,000*×*g for 60 minutes. Then, RNA from EVs was extracted as previously described. On the other hand, *in silico* reads were generated from different species (including human, bacteria, fungi and viruses) in different proportions (47.5M for human and between 25000 and 375000 for the rest of the species) *InSilicoSeq* v1.6.0 [19] (Table S3).

### Processing of taxa profiling data

For each sample, the raw profiling data table contains the genera and the reads assigned to each genus and profiler. The number of reads was corrected by a factor consisting of the total number of read pairs in the FASTQ of the sample divided by the median of the number of reads in the FASTQs of all samples.

The normalized number of reads was calculated by multiplying the raw number of reads by the inverse of this factor.

If *r*_*s,g*_ is the number of reads assigned to a sample *s* and a genus *g*, and *R*_*s*_ is the total number of reads in the FASTQ of sample *s*, the normalized count is 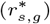:

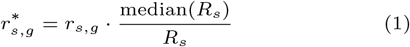

The mean value, standard deviation and coefficient of variation (CV) for all profilers were calculated from these normalized values.

For this analysis, we considered all samples as well as the bacterial controls.

To select relevant genera, the following flag system was applied: (a) if the genus contains information from only 1 profiler, 2 flags were added, and if it was from 2 profilers, 1 flag was added; (b) if the CV is greater than 1, 1 flag was added; (c) if the minimum number of reads is less than 100, 1 flag was added; and (d) if the mean number of reads across all profilers is less than 250, 1 flag was added. Then, the genera with only 0 or 1 flags were selected. The tables with information about genera selected before and after applying the flag system are available at Supplementary materials files *stats raw*.*xlsx* and *stats cut*.*xlsx*.

Finally, to make comparisons between samples, a common table is calculated with the mean across selected genera and samples, and genera that do not occur in more than 35% of the samples were removed. In addition, genera for which the log10 difference between the mean across samples and the mean across controls is less than 1 were discarded. This restriction is used to select only genera that have a significantly higher normalized frequency in the samples compared to the controls.

### Statistical analysis of samples across conditions

Based on the merged table of mean counts across samples, to perform statistical comparisons (e.g. HC vs RR-MS, sex) in all the contrasts proposed, a two-sided Mann-Whitney U test is computed with cutoff values of *α <* 0.05.

### **In silico** analysis

To evaluate the efficiency of the profilers with the artificially generated reads, a table was created with the reads assigned to each genus across the profilers. The efficiency was then measured depending on whether the genus to which the reads were assigned was included in the list of selected genera or not.

For the originally selected genera, the efficiency of a genus *g* and a profiler *p* was measured as follows

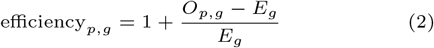

Where *O*_*p,g*_ is the number of reads the profiler *p* has assigned to genus *g* and *E*_*g*_ is the number of actual reads artificially generated for the genus.

On the other hand, the efficiency for genera to which no artificial reads have been assigned was measured as follows

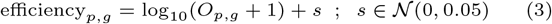

The random value *s* is only assigned for the representative function in order to avoid a collapse of several zero values.

## Results

### Analysis of profiler efficiency using **in silico** samples

This work aims to analyze the genera corresponding to the RNA associated with bacterial EVs in total plasma EVs from MS patients and HCs. To check the accuracy and bias of individual profilers, we generated *in silico* data consisting of random reads of selected genera and checked (1) how many of the actual reads were assigned to each genus and (2) how many reads were assigned to other genera that were not present in the selection sample. The results of this analysis are shown in Figure 2.

**Fig. 2.**
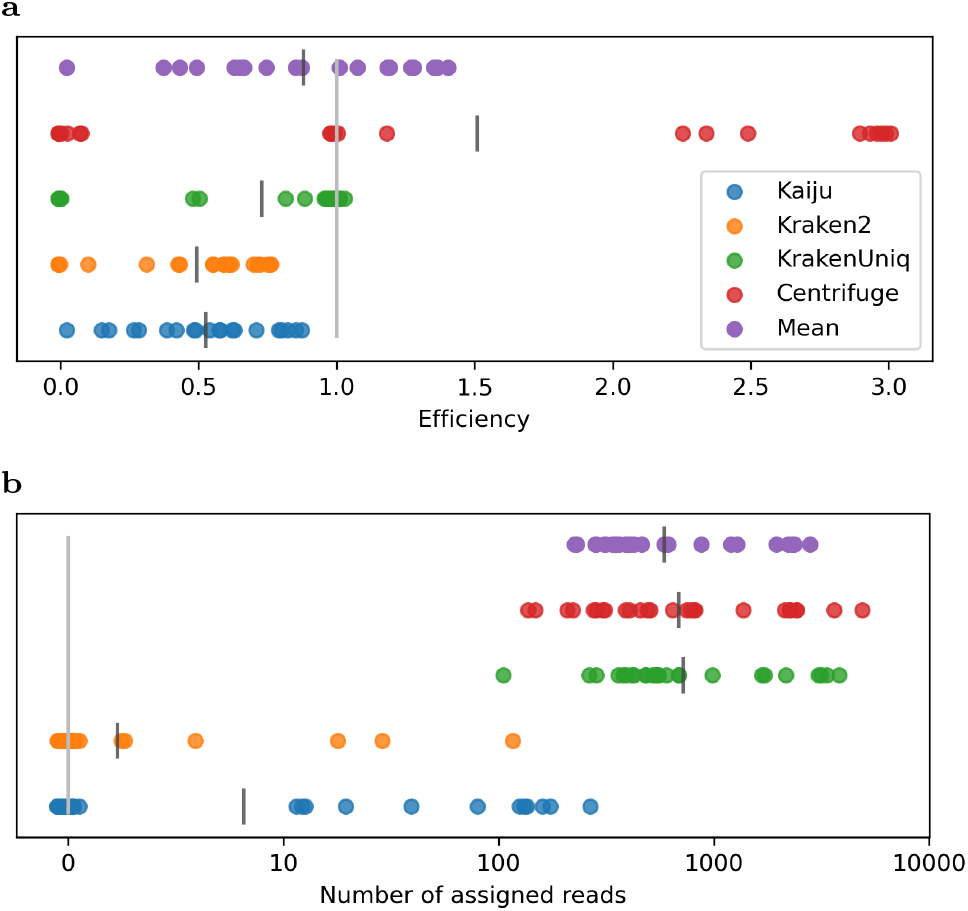
Efficiency of profilers with *in silico* samples for (a) genera present in the list of selected genera and (b) for genera absent from the list of selected genera. For panel (a) values closer to 1 represent a higher efficiency of the profiler, whereas for panel (b), a higher efficiency is achieved with values closer to 0 (gray bars). For each profiler, the black vertical bar represents the mean value across genera.

Focusing on the reads that were assigned to pre-existing genera, we find that *kaiju* and *kraken2* perform similarly, assigning on average half of the corresponding reads to each genus; *krakenuniq* shows the highest similarity to the original number of reads, although there are multiple genera (e.g. *Plenodomus, Alternatia, Burzaovirus*) that were not recognized, lowering the overall average; and centrifuge generally recognizes the reads in a much higher proportion (2-3 times), although for some genera (e.g. *Dietzia, Burzaovirus, Nickievirus*) it does not recognize the reads either. Interestingly, due to the overrepresentation of *centrifuge* and the underrepresentation of *kraken2* and *kaiju*, the mean values approached are remarkably close to the expected values. These values generally also match the percentage of reads assigned to non-human genera (see table S4). The percentages of assigned reads are 2.61% for *kaiju*, 2.63% for *kraken2*, 3.90% for *krakenuniq* and 8.14% for *centrifuge*. Considering that the expected percentage, i.e. the number of *in-silico* reads assigned non-human species, is 5%, we see that *kaiju* and *kraken2* represent almost half of the reads, centrifuge almost doubles and *krakenuniq* is the most accurate profiler.

Regarding the reads that were not assigned to selected genera, we note that *kaiju* and *kraken2* (especially *kraken2*) have extremely low amounts of misassigned reads. On the other hand, *krakenuniq* and *centrifuge* have higher assignment rates, with average values of almost 1,000 reads per genus. However, considering that the minimum original number of reads assigned to a genus is 25,000, the rate of misassignments is 1 to 2 orders of magnitude below the original number of reads and, thus, not alarming.

Therefore, each profiler has specific trade-offs regarding its functionality: *kraken2* and *kaiju* are more specific but have lower sensitivity. On the other hand, *krakenuniq* is more sensitive at the expense of a higher rate of reads assigned to other genera. Lastly, *centrifuge* is characterized by neither sensitivity nor specificity, but seems to complement the lower sensitivity values of *kraken2* and *kaiju*. Therefore, combining multiple profilers can increase the accuracy of predicting reads that are already assigned to existing genera while decreasing spurious assignment to other genera.

Finally, the set of mapped reads could strongly influence the choice of confidence parameters (e.g. hll-precision in *rkakenuniq* or e-value in *kaiju*). The reason for this is that the percentage of reads that need to be mapped after the 2nd mapping in the *in silico* dataset is 10.33%, which means that most non-human reads are probably not yet mapped and thus the strictness directly affects the proportion of mapped reads. We chose more stringent parameters to ensure that the selected genera were more likely to be biologically representative in the sample. However, a prior step of parameter tuning could be performed by adjusting the number of mapped reads to the expected number based on *in silico* datasets.

### The 3-fold human mapping and flag system are necessary for proper genera selection

A classic problem in taxa profiling studies is the correct assignment of reads to the corresponding taxa. It often happens that “contaminating” reads (i.e. reads originating from the host species, kit, etc.) are incorrectly assigned to other species, leading to a high false positive assignment rate [11]. To minimize this problem, we have included two important steps in the mapping and processing pipeline: 3-fold mapping to the human genome and the flag system.

The 3-fold mapping consists of two mapping steps using the human genomes GRCh38 and CHM13 with two different aligners (STAR and *bowtie2*) and mapping to the human genome in all profilers. In this analysis, we find that for MS samples, a high percentage of reads (*>* 90%) are assigned to the human genome only in the first map. Strikingly, however, an additional 1-3% of reads (about 0.5-1.5 million) are assigned in the second map. In terms of mapping to non-human genera, we see that the third map filters out an additional 1-3% of reads in all samples, although the results vary greatly depending on the profiler. Similarly for the control samples analyzed with *L. acidophilus* and *B. lactis*, we find that for *L. acidophilus* almost 4% of the reads are assigned to the human genome. This effect is likely caused by cross-contamination during sample handling or due to computational artifacts of reads mapping to ortholog areas of the human genome. Regardless of the cause, this scenario shows the 3-step host mapping to be highly relevant.

Based on *in silico* data, and considering that only 5% of the reads belong to non-human genera, another 5% of the reads belonging to humans cannot be assigned to the GRCh38 and CHM13 genomes, emphasizing the need to use a human reference in the profiler database of genomes when analyzing other species. In some cases, such as *kaiju* or *kraken2*, the lower assignment rate to non-human genera may be due to the fact that many reads may have a high homology rate with human orthologs.

The next step in the processing pipeline is to assign flags to ensure that only genera that are consistently assigned by most profilers and that have a minimum number of reads per sample are selected, thereby minimizing the risk of false-positive bias. When considering the genera assigned in the *in silico* dataset, which consisted of 20 non-human genera, 47 genera passed the flag system thresholds (the total number of genera initially assigned was 936). However, the number of reads assigned to the expected 20 genera was three orders of magnitude higher than for the 27 non-original genera (mean number of reads across genera and profilers: 112608 vs. 836), as shown in Figure S1. In addition, most of these genera were assigned a much higher number of reads by *krakenuniq* and *centrifuge*, which are biased to be less sensitive (Figure 2b).

Using control samples, 602 and 189 genera are identified in the raw results of *L. acidophilus* and *B. lactis* respectively. After applying the flag system, however, the assignment is reduced to only 50 and 14 genera, respectively. Most of the non-controlled genera that are not assigned to humans, such as *Cutibacterium* or *Enterobacter*, are described as part of the human microbiome [20, 21], so their presence in the analysis is probably due to contamination during sample manipulation.

Although we are aware of the limitations of the 3-way mapping and flagging system, namely the higher rate of false negatives, it shows a drastic improvement in the results obtained in *in silico* and in control samples, and we believe that this system is necessary to ensure a more accurate putative characterization of the genera present in the biological samples from human donors.

## Relevance of normalisation to remove contaminant genera

Another important step in the processing of samples is normalization. In our work, we opted for the median normalization step, where the median of the total number of reads across all samples (including bacterial controls) is set as the key value. In our data, the total number of reads of total plasma EVs from MS samples was about 25 times higher than bacterial EVs of control samples (*∼*50M reads vs. *∼*2M reads). Therefore, without normalization, the number of reads assigned to specific genera might be underrepresented.

Figure 3 shows the genera for which the number of reads assigned to total plasma EVs from MS samples is at least 10 times higher than for the bacterial EVs technical control samples. We observe several genera such as *Dietzia, Nesteronkonia* or *Mitsuaria* that have no counts in bacterial control samples, while others such as *Sphingomonas, Acidovorax, Acinetobacter* or *Corynebacterium* have a variable number of reads.

**Fig. 3.**
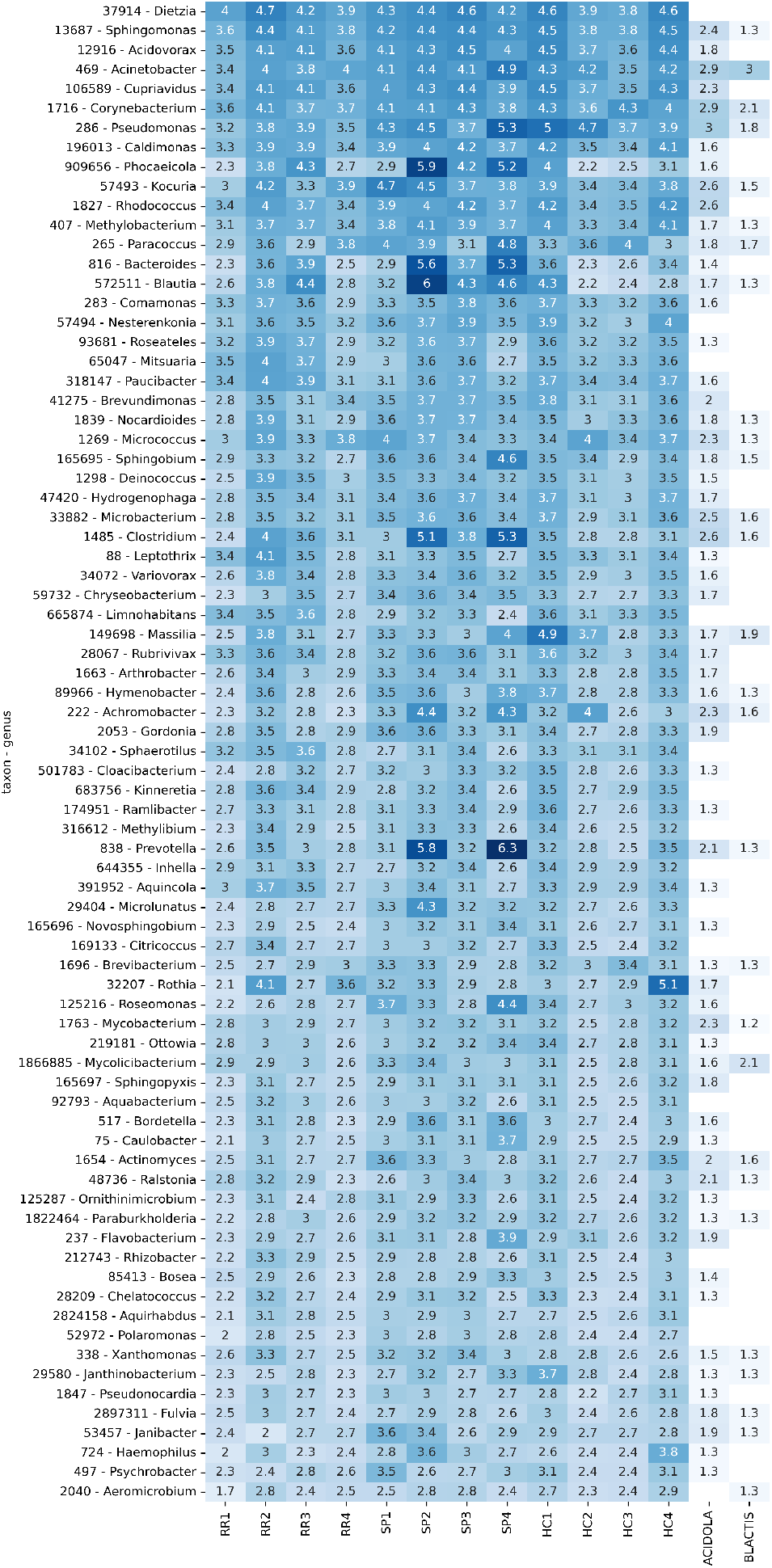
Heatmap showing genera that pass the cut criteria of being 10 times more abundant in samples than in controls, with normalisation.

Interestingly, a comparison of the number and type of genera that failed the cutoff of being 10 times more abundant than in control samples, with and without normalization, (Figures S2 and S4) shows that the number of discarded genera is higher after normalization (24 vs. 5). Interestingly, genera such as *Cutibacterium, Staphylococcus* or *Dermacoccus*, which did not make the cut, were characterized as skin commensals [20, 21] and are therefore likely contaminants in the samples. This shows how important it is to carry out controls to discard putative contaminant genera from the samples.

## Differential abundance of genera across conditions

After obtaining the list of genera highly likely to be present in the samples and not significantly represented in the controls, we performed several comparisons between MS types and HC. The species with differential abundance between the comparisons are shown in Figure S5.

When comparing RR-MS and HC, only the genus *Janibacter* appears to have a differential abundance (more abundant in HC samples), although the size effect is not remarkable (log2FC: 0.92, *p*-value:0.029). Regarding SP-MS vs. HC, three genera show a statistically significant differential abundance in SP-MS samples: *Xanthomonas, Chryseobacterium* and *Caulobacter* ; all three more abundant in SP-MS, with a *p*-value of 0.029 and log2FC of 1.69, 1.21 and 1.81, respectively.

Finally, the comparison between SP-MS and RR-MS revealed 5 statistically significant genera (*p*-value: 0.029), 3 of which were more abundant in SP-MS and had relevant log2FC values: *Microlunatus* (3.82), *Sphingobium* (3.48) and *Novosphingobium* (2.00).

The comparison between the sexes across all conditions did not reveal any statistically significant differentially abundant genus. Similarly, a comparison between combined MS samples (RR-MS and SP-MS) did not yield any statistically significant differentially abundant genus. Lastly, it should be noted that no *p*-value reached statistical significance after correction for multiple comparisons.

## Discussion

The role of microorganisms in the human body is of great significance, as the body is composed of a similar number of bacterial and human cells [22]. Therefore, it is crucial to further study human-human, human-bacterial and bacterial-bacterial communication, both in maintaining homeostasis and in the context of disease. One approach to this is to detect microbial reads in the NGS studies in human tissues, but it is essential to carefully retrieve, process, and analyze samples to avoid false positive bias and minimize the risk of error.

Recently, a reanalysis by Gihawi et al. of a study that reported strong correlations between DNA signatures of microbial organisms and 33 different cancer types [22] has noted this effect. The authors of the reanalysis claimed that the use of lenient thresholds and the lack of thorough mapping to the host genome led to an increase in the detection of novel bacterial species, including some belonging to the category of extremophiles which, thus, were highly unlikely to be found in human cancer samples.

In our study, we support the findings of Gihawi et al. by demonstrating the need for a robust pipeline of mapping to the host genome to reduce the rate of false positive assignment of reads to microbial species. Additionally, we recommend using multiple profilers and a flag system to minimize non-redundant results. We also emphasize the importance of using control samples, both *in silico* and *in vitro*, to monitor sampling and processing biases as much as possible. Our results demonstrate, that with adequate caution, Bacterial reads can be found when analyzing by RNAseq the cargo of EVs in human plasma.

The analysis identified promising candidates for further investigation within both the MS and control groups. This is particularly significant due to previously documented shifts in the stool microbiome of MS patients [23, 9]. Of note, a comparison of the genera described in this work with species reported by these studies shows no presence of statistically significant genera in the reported results. Interestingly, both *Blautia* and *Clostridium* show a relatively increased abundance in MS samples compared to HC (log2FC of 0.44 and 0.32, *p*-values of 0.07 and 0.21), whereas [9] report a decrease in MS samples. It should be noted that these results cannot be directly comparable due to the differences in sample type (stool samples vs EVs) and analysis type (16S rRNAvs RNAseq). In this analysis we are not characterizing the diversity of the bacterial kingdom through DNA-sequencing but studying the messages that the bacterial deliver through the putative bEVS to communicate with other tissues. Although we can find the reads, more deep in the NGS is needed to study the specific genes and their potential functions.

Several genera were consistently present across all samples and have been detected in other human-associated environments. For example, *Dietzia* has been isolated from blood in sepsis and skin conditions [24], and its putative potential as a probiotic for Crohn’s disease after being tested in cows with a disease model has been explored [25]. Similarly, *Nesterenkonia* and *Sphingomonas* have been documented in faecal and skin samples [26, 27]. *Acidovorax* has been associated with stroke patients [28]. Lastly, *Cupriavidus*, known for its metal resistance [29], has been found in hospital sink traps [30] and is capable of causing infections [31, 32, 33].

Regarding genera differentially abundant between conditions, some of them are worth mentioning. *Janibacter*, more abundant in SP-MS than in HCs, has been positively associated with increased Parkinson’s Disease Rating Scale values [34]. *Xanthomonas* and *Chryseobacterium*, two genera with more representation in RR-MS than in controls, have been isolated from tumoral lesions [35] and in urinary infections [36, 37] respectively.

Lastly, regarding genera more abundant in SP-MS than in RR-MS, three are of interest. Firstly, *Sphingobium* has been discovered in patients undergoing peritoneal dialysis [38]. *Novosphingobium* is associated with colorectal cancer [39], chronic obstructive pulmonary disease (COPD) [40], and biliary cirrhosis [41]. Lastly, *Microlunatus*, despite being differentially abundant in our study samples, no presence in human has been previously recorded, although it has been characterized from soil samples [42].

It is noteworthy that many of these identified genera are not human-specific and have been found in plants and food sources (e.g. *Sphingobium* [43], *Xanthomonas* [44], *Acidovorax* [45], or *Microlunatus* [42]. This suggests that some amplified genetic material from EVs may originate from ingested, breathed or, more generally, exogenously introduced species with subsequent transport through the bloodstream as part of the EV cargo or the widely reported EV-corona [46]. This finding highlights the potential role of diet and, more generally, the environment, in influencing the microbiome in MS, given the established link between diet and gut microbiota composition [47, 48, 49].

To summarise, based on these results, the study proposed in this article aims to contribute to the reduction of false positive bias in taxa profiling originating from EVs or alternative sources, as well as enhancing the replicability and robustness of future research employing such methods.

### Study limitations

Despite the relevance of the results from this study, several limitations must be addressed. Although the idea of the project is a proof of concept for the analysis tools, the results of the DE genera must be taken with caution due to the size of the groups. Bigger series must be performed to produce reliable DE lists. Additionally, these results, but not the analysis flow proposed, are dependent on the methodology used to isolate the EVs, a common issue for all EV studies. Lastly, it should be considered that part of the retrieved bacterial reads could not be directly associated with the presence of bEVs, but rather from other forms of genetic material including RNA derived from blood-circulating bacteria.

## Supporting information

CSV and XLSX files obtained during data analysis

## Data availability

The data underlying this article are available in GEO database and can be accessed with the following accession number: GSE255317. All the analyses described in the methods can be accessed at the GitHub repository https://github.com/NanoNeuro/EV_taxprofiling.

## Acknowledgements

The authors want to acknowledge all the patients who donated samples for this study.

## References

1. Ainhoa Alberro, Leire Iparraguirre, Adelaide Fernandes, and David Otaegui. Extracellular vesicles in blood: Sources, effects, and applications. International Journal of Molecular Sciences, 22(15):8163, July 2021.

2. Masanori Toyofuku, Stefan Schild, Maria Kaparakis-Liaskos, and Leo Eberl. Composition and functions of bacterial membrane vesicles. Nature Reviews Microbiology, 21(7):415–430, March 2023.

3. Clement Yaw Effah, Xianfei Ding, Emmanuel Kwateng Drokow, Xiang Li, Ran Tong, and Tongwen Sun. Bacteria-derived extracellular vesicles: endogenous roles, therapeutic potentials and their biomimetics for the treatment and prevention of sepsis. Frontiers in Immunology, 15, February 2024.

4. Niloufar Hosseini-Giv, Alyza Basas, Chloe Hicks, Emad El-Omar, Fatima El-Assaad, and Elham Hosseini-Beheshti. Bacterial extracellular vesicles and their novel therapeutic applications in health and cancer. Frontiers in Cellular and Infection Microbiology, 12, November 2022.

5. Katja Koeppen, Thomas H. Hampton, Michael Jarek, Maren Scharfe, Scott A. Gerber, Daniel W. Mielcarz, Elora G. Demers, Emily L. Dolben, John H. Hammond, Deborah A. Hogan, and Bruce A. Stanton. A novel mechanism of host-pathogen interaction through srna in bacterial outer membrane vesicles. PLOS Pathogens, 12(6):e1005672, June 2016.

6. Natayme R. Tartaglia, Koen Breyne, Evelyne Meyer, Chantal Cauty, Julien Jardin, Denis Chrétien, Aurélien Dupont, Kristel Demeyere, Nadia Berkova, Vasco Azevedo, Eric Guédon, and Yves Le Loir. Staphylococcus aureus extracellular vesicles elicit an immunostimulatory response in vivo on the murine mammary gland. Frontiers in Cellular and Infection Microbiology, 8, August 2018.

7. Kerstin Berer, Lisa Ann Gerdes, Egle Cekanaviciute, Xiaoming Jia, Liang Xiao, Zhongkui Xia, Chuan Liu, Luisa Klotz, Uta Stauffer, Sergio E. Baranzini, Tania Kümpfel, Reinhard Hohlfeld, Gurumoorthy Krishnamoorthy, and Hartmut Wekerle. Gut microbiota from multiple sclerosis patients enables spontaneous autoimmune encephalomyelitis in mice. Proceedings of the National Academy of Sciences, 114(40):10719–10724, September 2017.

8. Egle Cekanaviciute, Bryan B. Yoo, Tessel F. Runia, Justine W. Debelius, Sneha Singh, Charlotte A. Nelson, Rachel Kanner, Yadira Bencosme, Yun Kyung Lee, Stephen L. Hauser, Elizabeth Crabtree-Hartman, Ilana Katz Sand, Mar Gacias, Yunjiao Zhu, Patrizia Casaccia, Bruce A. C. Cree, Rob Knight, Sarkis K. Mazmanian, and Sergio E. Baranzini. Gut bacteria from multiple sclerosis patients modulate human t cells and exacerbate symptoms in mouse models. Proceedings of the National Academy of Sciences, 114(40):10713–10718, September 2017.

9. Xiaoyuan Zhou, Ryan Baumann, Xiaohui Gao, Myra Mendoza, Sneha Singh, Ilana Katz Sand, Zongqi Xia, Laura M. Cox, Tanuja Chitnis, Hongsup Yoon, Laura Moles, Stacy J. Caillier, Adam Santaniello, Gail Ackermann, Adil Harroud, Robin Lincoln, Refujia Gomez, Antonio González Penã, Elise Digga, Daniel Joseph Hakim, Yoshiki Vazquez-Baeza, Karthik Soman, Shannon Warto, Greg Humphrey, Mauricio Farez, Lisa Ann Gerdes, Jorge R. Oksenberg, Scott S. Zamvil, Siddharthan Chandran, Peter Connick, David Otaegui, Tamara Castillo-Trivinõ, Stephen L. Hauser, Jeffrey M. Gelfand, Howard L. Weiner, Reinhard Hohlfeld, Hartmut Wekerle, Jennifer Graves, Amit Bar-Or, Bruce A.C. Cree, Jorge Correale, Rob Knight, and Sergio E. Baranzini. Gut microbiome of multiple sclerosis patients and paired household healthy controls reveal associations with disease risk and course. Cell, 185(19):3467–3486.e16, September 2022.

10. Leire Iparraguirre, Ainhoa Alberro, Thomas B. Hansen, Tamara Castillo-Trivinõ, Maider Munõz-Culla, and David Otaegui. Profiling of plasma extracellular vesicle transcriptome reveals that circrnas are prevalent and differ between multiple sclerosis patients and healthy controls. Biomedicines, 9(12):1850, December 2021.

11. Abraham Gihawi, Yuchen Ge, Jennifer Lu, Daniela Puiu, Amanda Xu, Colin S. Cooper, Daniel S. Brewer, Mihaela Pertea, and Steven L. Salzberg. Major data analysis errors invalidate cancer microbiome findings. mBio, 14(5), October 2023.

12. Philip A. Ewels, Alexander Peltzer, Sven Fillinger, Harshil Patel, Johannes Alneberg, Andreas Wilm, Maxime Ulysse Garcia, Paolo Di Tommaso, and Sven Nahnsen. The nf-core framework for community-curated bioinformatics pipelines. Nature Biotechnology, 38(3):276–278, February 2020.

13. Alexander Dobin, Carrie A. Davis, Felix Schlesinger, Jorg Drenkow, Chris Zaleski, Sonali Jha, Philippe Batut, Mark Chaisson, and Thomas R. Gingeras. Star: ultrafast universal rna-seq aligner. Bioinformatics, 29(1):15–21, October 2012.

14. Peter Menzel, Kim Lee Ng, and Anders Krogh. Fast and sensitive taxonomic classification for metagenomics with kaiju. Nature Communications, 7(1), April 2016.

15. Derrick E. Wood, Jennifer Lu, and Ben Langmead. Improved metagenomic analysis with kraken 2. Genome Biology, 20(1), November 2019.

16. F. P. Breitwieser, D. N. Baker, and S. L. Salzberg. Krakenuniq: confident and fast metagenomics classification using unique k-mer counts. Genome Biology, 19(1), November 2018.

17. Daehwan Kim, Li Song, Florian P. Breitwieser, and Steven L. Salzberg. Centrifuge: rapid and sensitive classification of metagenomic sequences. Genome Research, 26(12):1721–1729, October 2016.

18. Moritz E. Beber, Maxime Borry, Sofia Stamouli, and James A. Fellows Yates. Taxpasta: Taxonomic profile aggregation and standardisation. Journal of Open Source Software, 8(87):5627, July 2023.

19. Hadrien Gourlé, Oskar Karlsson-Lindsjö, Juliette Hayer, and Erik Bongcam-Rudloff. Simulating illumina metagenomic data with insilicoseq. Bioinformatics, 35(3):521–522, July 2018.

20. Clio Dessinioti and Andreas Katsambas. The microbiome and acne: Perspectives for treatment. Dermatology and Therapy, January 2024.

21. Jonathan N. V. Martinson, Nicholas V. Pinkham, Garrett W. Peters, Hanbyul Cho, Jeremy Heng, Mychiel Rauch, Susan C. Broadaway, and Seth T. Walk. Rethinking gut microbiome residency and the enterobacteriaceae in healthy human adults. The ISME Journal, 13(9):2306–2318, May 2019.

22. Ron Sender, Shai Fuchs, and Ron Milo. Revised estimates for the number of human and bacteria cells in the body. PLOS Biology, 14(8):e1002533, August 2016.

23. Florence Thirion, Finn Sellebjerg, Yong Fan, Liwei Lyu, Tue H. Hansen, Nicolas Pons, Florence Levenez, Benoit Quinquis, Evelina Stankevic, Helle B. Søndergaard, Thomas M. Dantoft, Casper S. Poulsen, Sofia K. Forslund, Henrik Vestergaard, Torben Hansen, Susanne Brix, Annette Oturai, Per Soelberg Sørensen, Stanislav D. Ehrlich, and Oluf Pedersen. The gut microbiota in multiple sclerosis varies with disease activity. Genome Medicine, 15(1), January 2023.

24. Roland J. Koerner, Michael Goodfellow, and Amanda L. Jones. The genusdietzia: a new home for some known and emerging opportunist pathogens. FEMS Immunology & Medical Microbiology, 55(3):296–305, April 2009.

25. Robert E. Click and Craig L. Van Kampen. Assessment of dietzia subsp.c79793-74 for treatment of cattle with evidence of paratuberculosis. Virulence, 1(3):145–155, May 2010.

26. Sophie Edouard, Senthil Sankar, Nicole Prisca Makaya Dangui, Jean-Christophe Lagier, Caroline Michelle, Didier Raoult, and Pierre-Edouard Fournier. Genome sequence and description of nesterenkonia massiliensis sp. nov. strain np1t. Standards in Genomic Sciences, 9(3):866–882, April 2014.

27. Izabela Szczerba. [occurrence and number of bacteria from the micrococcus, kocuria, nesterenkonia, kytococcus and dermacoccus genera on skin and mucous membranes in humans]. Medycyna doswiadczalna i mikrobiologia, 55(1):67—74, 2003.

28. Yaning Zang, Xigui Lai, Conghui Li, Dongfang Ding, Ying Wang, and Yi Zhu. The role of gut microbiota in various neurological and psychiatric disorders—an evidence mapping based on quantified evidence. Mediators of Inflammation, 2023:1–16, February 2023.

29. Rob Van Houdt, Ann Provoost, Ado Van Assche, Natalie Leys, Bart Lievens, Kristel Mijnendonckx, and Pieter Monsieurs. Cupriavidus metallidurans strains with different mobilomes and from distinct environments have comparable phenomes. Genes, 9(10):507, October 2018.

30. James Butler, Sean D. Kelly, Katie J. Muddiman, Alexandros Besinis, and Mathew Upton. Hospital sink traps as a potential source of the emerging multidrug-resistant pathogen cupriavidus pauculus: characterization and draft genome sequence of strain mf1. Journal of Medical Microbiology, 71(2), February 2022.

31. Zhen Zhang, Wanyan Deng, Shuling Wang, Lanlan Xu, Ling Yan, and Pu Liao. First case report of infection caused by cupriavidus gilardii in a non-immunocompromised chinese patient. IDCases, 10:127–129, 2017.

32. Stéphanie Langevin, Jean Vincelette, Sadjia Bekal, and Christiane Gaudreau. First case of invasive human infection caused by cupriavidus metallidurans. Journal of Clinical Microbiology, 49(2):744–745, February 2011.

33. Daniel Tena, Cristina Losa, María José Medina, and Juan Antonio Sáez-Nieto. Muscular abscess caused by cupriavidus gilardii in a renal transplant recipient. Diagnostic Microbiology and Infectious Disease, 79(1):108–110, May 2014.

34. Yiwei Qian, Xiaodong Yang, Shaoqing Xu, Chunyan Wu, Nan Qin, Sheng-Di Chen, and Qin Xiao. Detection of microbial 16s rrna gene in the blood of patients with parkinson’s disease. Frontiers in Aging Neuroscience, 10, May 2018.

35. William F. Marshall, Michael R. Keating, John P. Anhalt, and James M. Steckelberg. Xanthomonas maltophilia: An emerging nosocomial pathogen. Mayo Clinic Proceedings, 64(9):1097–1104, September 1989.

36. M. Isabel Acosta-Ochoa, Antonio Rodrigo-Parra, Flor Rodríguez-Martín, and Antonio Molina-Miguel. Infección urinaria por chryseobacterium indologenes. Nefrología, (33), July 2013.

37. Dora E Izaguirre-Anariba and Vel Sivapalan. Chryseobacterium indologenes, an emerging bacteria: A case report and review of literature. Cureus, January 2020.

38. Sujith K Palleti, Santhoshi R Bavi, Margaret Fitzpatrick, and Anuradha Wadhwa. First case report of sphingobium lactosutens as a human pathogen causing peritoneal dialysis-related peritonitis. Cureus, July 2022.

39. Bin Zhou, Linli Shi, Min Jin, Mingxia Cheng, Dandan Yu, Lei Zhao, Jieying Zhang, Yu Chang, Tao Zhang, and Hongli Liu. Caulobacter and novosphingobium in tumor tissues are associated with colorectal cancer outcomes. Frontiers in Oncology, 12, January 2023.

40. Alleluiah Rutebemberwa, Mark J. Stevens, Mario J. Perez, Lynelle P. Smith, Linda Sanders, Gregory Cosgrove, Charles E. Robertson, Rubin M. Tuder, and J. Kirk Harris. Novosphingobium and its potential role in chronic obstructive pulmonary diseases: Insights from microbiome studies. PLoS ONE, 9(10):e111150, October 2014.

41. Marshall M. Kaplan. Novosphingobium aromaticivorans: A potential initiator of primary biliary cirrhosis. The American Journal of Gastroenterology, 99(11):2147–2149, November 2004.

42. K. Nakamura, A. Hiraishi, Y. Yoshimi, M. Kawaharasaki, K. Masuda, and Y. Kamagata. Microlunatus phosphovorus gen. nov., sp. nov., a new gram-positive polyphosphate-accumulating bacterium isolated from activated sludge. International Journal of Systematic Bacteriology, 45(1):17–22, January 1995.

43. Yang Jia, Adel Eltoukhy, Junhuan Wang, Xianjun Li, Thet Su Hlaing, Mar Mar Aung, May Thet Nwe, Imane Lamraoui, and Yanchun Yan. Biodegradation of bisphenol a by sphingobium sp. yc-jy1 and the essential role of cytochrome p450 monooxygenase. International Journal of Molecular Sciences, 21(10):3588, May 2020.

44. Shi-Qi An, Neha Potnis, Max Dow, Frank-Jörg Vorhölter, Yong-Qiang He, Anke Becker, Doron Teper, Yi Li, Nian Wang, Leonidas Bleris, and Ji-Liang Tang. Mechanistic insights into host adaptation, virulence and epidemiology of the phytopathogenxanthomonas. FEMS Microbiology Reviews, 44(1):1–32, October 2019.

45. D.L. Hopkins and C.M. Thompson. Seed transmission of acidovorax avenae subsp. citrulli in cucurbits. HortScience, 37(6):924–926, October 2002.

46. Eszter Á. Tóth, Lilla Turiák, Tamás Visnovitz, Csaba Cserép, Anett Mázló, Barbara W. Sódar, András I. Försönits, Gábor Pet?ovári, Anna Sebestyén, Zsolt Komlósi, László Drahos, Ágnes Kittel, György Nagy, Attila Bácsi, Ádám Dénes, Yong Song Gho, Katalin É. Szabó-Taylor, and Edit I. Buzás. Formation of a protein corona on the surface of extracellular vesicles in blood plasma. Journal of Extracellular Vesicles, 10(11), September 2021.

47. Laura A. Pace and Sheila E. Crowe. Complex relationships between food, diet, and the microbiome. Gastroenterology Clinics of North America, 45(2):253–265, June 2016.

48. Herbert Tilg and Alexander R. Moschen. Food, immunity, and the microbiome. Gastroenterology, 148(6):1107–1119, May 2015.

49. Noah Voreades, Anne Kozil, and Tiffany L. Weir. Diet and the development of the human intestinal microbiome. Frontiers in Microbiology, 5, September 2014.

