## Supplementary material for "A proposed workflow to analyze bacterial transcripts in RNAseq from blood extracellular vesicles of people with Multiple Sclerosis": CSV and XLSX files obtained during data analysis: README

artificial_species_table.csv - Information regarding artificially-generated species (kingdom, genus, species, NCBI Tax ID, relative abundance and number of reads).
comparison_conditions.xlsx - Table with data used to perform the comparisons for each condition and report their p-values. 
percentage_reads_profilers.csv - Number of reads, and relative percentage to total, assigned in each of the pipeline steps.
stats_cut.xlsx - Information about selected genera (reads assigned per profiler, normalised read counts, mean, std, CV and flags).
stats_raw.xlsx - Information about all genera (reads assigned per profiler, normalised read counts, mean, std, CV and flags).
