## Supplementary Material for "A proposed workflow to analyze bacterial transcripts in RNAseq from blood extracellular vesicles of people with Multiple Sclerosis"

### 1 Supplementary Tables

Table S1: Main clinical and demographical characteristics of individuals enrolled in the study classified by disease status.

| Disease status | Sex | Age | EDSS | Evol. time | AOO |
|---|---|---|---|---|---|
| RR-MS (n=10) | Male (n=5) | 39.7 (±13.7) | 2 (0-3.5) | 16.1 (±9.8) | 23.6 (±9.1) |
| | Female (n=5) | 42.0 (±19.8) | 0 (0-2) | 13.2 (±12.2) | 28.8 (±9.9) |
| SP-MS (n=10) | Male (n=5) | 54.8 (±6.2) | 6 (6-8.5) | 20.7 (±8.7) | 34.1 (±9.5) |
| | Female (n=5) | 48.7 (±6.7) | 7 (4-8) | 22.5 (±5.0) | 26.2 (±6.6) |
| HC (n=8) | Male (n=4) | 56.1 (±11.9) | - | - | - |
| | Female (n=4) | 53.6 (±11.8) | - | - | - |

Abbreviations: RR-MS, relapsing-remitting multiple sclerosis; SP-MS. Secondary-progressive multiple sclerosis; HC, healthy control; EDSS, Expanded Disability Status Scale; Evol. Time: Evolution time; AOO, Age of Onset. Age, evolution time and AOO data are presented as "average (standard deviation)", EDSS data are shown as "median (range)".

Table S2: Patient description of each pool.

| Pool name | Patient | Sample collection | Disease Status | Sex |
|---|---|---|---|---|
| Pool 1 | EVRR1 | 2/18/2010 | RR-MS | M |
| | EVRR2 | 2/25/2012 | RR-MS | M |
| | EVRR3 | 4/23/2009 | RR-MS | M |
| Pool 2 | EVRR4 | 2/25/2010 | RR-MS | M |
| | EVRR5 | 6/19/2013 | RR-MS | M |
| Pool 3 | EVRR6 | 12/2/2010 | RR-MS | F |
| | EVRR7 | 5/26/2009 | RR-MS | F |
| | EVRR8 | 7/9/2009 | RR-MS | F |
| Pool 4 | EVRR9 | 7/9/2009 | RR-MS | F |
| | EVRR10 | 9/16/2010 | RR-MS | F |
| Pool 5 | EVSP1 | 3/5/2009 | SP-MS | M |
| | EVSP2 | 5/28/2009 | SP-MS | M |
| | EVSP3 | 3/28/2011 | SP-MS | M |
| Pool 6 | EVSP4 | 12/23/2010 | SP-MS | M |
| | EVSP5 | 4/28/2009 | SP-MS | M |
| Pool 7 | EVSP6 | 2/17/2011 | SP-MS | F |
| | EVSP7 | 9/12/2000 | SP-MS | F |
| | EVSP8 | 9/28/2010 | SP-MS | F |
| Pool 8 | EVSP9 | 1/13/2011 | SP-MS | F |
| | EVSP10 | 10/7/2010 | SP-MS | F |
| Pool 9 | EVHC1 | 12/2/2004 | HC | M |
| | EVHC2 | 9/22/2009 | HC | M |
| Pool 10 | EVHC3 | 9/24/2009 | HC | M |
| | EVHC4 | 9/24/2009 | HC | M |
| Pool 11 | EVHC5 | 12/16/2009 | HC | F |
| | EVHC6 | 2/26/2009 | HC | F |
| Pool 12 | EVHC7 | 7/4/2011 | HC | F |
| | EVHC8 | 7/15/2011 | HC | F |

### 2 Supplementary figures

Table S3: Summary table of species used to generate the artificial reads.

| **Kingdom** | **Genus** | **Species** | **NCBI Tax ID** | **Rel. abundance** | **N reads** |
|---|---|---|---|---|---|
| Animal | Homo | sapiens | 9606 | 0.95 | 47500000 |
| Bacteria | Cutibacterium | acnes | 1747 | 0.0075 | 375000 |
| Bacteria | Dietzia | maris | 37915 | 0.005 | 250000 |
| Bacteria | Bifidobacterium | bifidum | 1681 | 0.005 | 250000 |
| Bacteria | Lactobacillus | acidophilus | 1579 | 0.0025 | 125000 |
| Bacteria | Bacteroides | fragilis | 817 | 0.0025 | 125000 |
| Bacteria | Roseburia | intestinalis | 166486 | 0.0025 | 125000 |
| Bacteria | Stenotrophomonas | maltophilia | 40324 | 0.0015 | 75000 |
| Bacteria | Sphingomonas | paucimobilis | 13689 | 0.0015 | 75000 |
| Bacteria | Taylorella | equigenitalis | 29575 | 0.001 | 50000 |
| Bacteria | Kinneretia | asaccharophila | 582607 | 0.0005 | 25000 |
| Fungi | Aspergillus | chevalieri | 182096 | 0.005 | 250000 |
| Fungi | Malassezia | restricta | 76775 | 0.0025 | 125000 |
| Fungi | Saccharomyces | cerevisiae | 4932 | 0.0025 | 125000 |
| Fungi | Plenodomus | lingam | 5022 | 0.0015 | 75000 |
| Fungi | Alternaria | alternata | 5599 | 0.0015 | 75000 |
| Virus | Burzaovirus | intestinihominis | 2955562 | 0.0025 | 125000 |
| Virus | Elvirus | EL | 1984787 | 0.0015 | 75000 |
| Virus | Lentivirus | HIV1 | 11676 | 0.0015 | 75000 |
| Virus | Nickievirus | nickie | 2560688 | 0.0015 | 75000 |
| Virus | Lymphocryptovirus | humangamma4 | 3050299 | 0.0005 | 25000 |

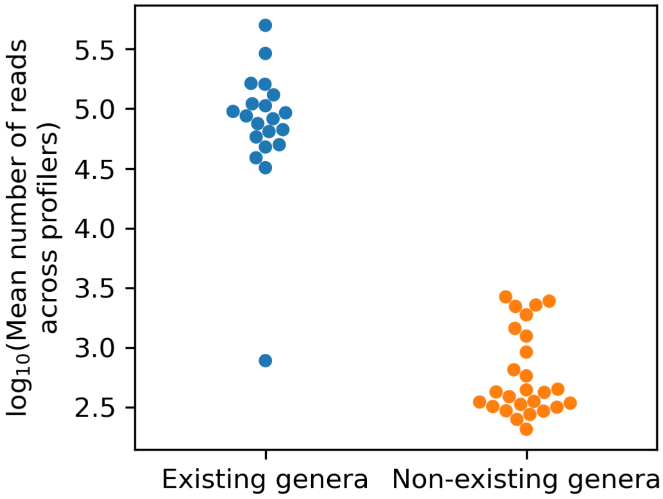


Figure S1: Swarmplot showing the mean number of reads assigned by each profiler to each genera in the *in silico* dataset. Genera are separated by whether they were originally included in the dataset FASTQ file, or not.

Table S4: Sample information and percentages of reads used in the next step of the processing pipeline. For instance, POOL1 sample originally contained 56.55 million of reads (100%). Of that amount, 6.67% (3.77M) were retained after the first map (the rest were mapped to human); and 5.81% (3.28M) were retained after the second map. Lastly of these reads, 5.50% (3.11M) were mapped to genera by *kraken2*, and 0.38% (0.21M) were mapped to non-human genera.
Column labels: MS (multiple sclerosis type), KAI (*kaiju*), KR2 (*kraken2*), KRU (*krakenuniq)*, CEN (*centrifuge)*.

| | | | | | | **WITH HUMAN READS** | | | | **WITHOUT HUMAN READS** | | | |
|---|---|---|---|---|---|---|---|---|---|---|---|---|---|
| | **MS** | **Sex** | **Base (M)** | **1st map** | **2nd map** | **KAI** | **KR2** | **KRU** | **CEN** | **KAI** | **KR2** | **KRU** | **CEN** |
| **POOL1** | RR | M | 56.55 | 6.67 | 5.81 | 0.92 | 5.50 | 5.45 | 3.78 | 0.05 | 0.38 | 0.36 | 0.67 |
| **POOL2** | RR | M | 42.21 | 11.65 | 10.59 | 1.24 | 8.39 | 7.98 | 6.51 | 0.29 | 2.89 | 2.53 | 2.37 |
| **POOL3** | RR | F | 48.79 | 9.31 | 6.79 | 0.51 | 5.39 | 5.24 | 3.68 | 0.08 | 1.43 | 1.28 | 1.34 |
| **POOL4** | RR | F | 47.15 | 9.09 | 6.13 | 0.56 | 5.31 | 5.25 | 3.62 | 0.06 | 0.66 | 0.59 | 0.78 |
| **POOL5** | SP | M | 44.69 | 7.93 | 6.22 | 0.46 | 4.79 | 4.65 | 3.45 | 0.12 | 1.63 | 1.52 | 1.49 |
| **POOL6** | SP | M | 55.64 | 43.75 | 42.77 | 1.10 | 32.72 | 27.95 | 24.54 | 0.90 | 31.03 | 26.27 | 23.36 |
| **POOL7** | SP | F | 55.70 | 9.30 | 7.28 | 0.45 | 5.29 | 5.02 | 4.08 | 0.15 | 2.23 | 1.96 | 1.79 |
| **POOL8** | SP | F | 47.83 | 29.66 | 28.57 | 0.76 | 23.83 | 23.64 | 18.15 | 0.40 | 20.69 | 20.51 | 16.08 |
| **POOL9** | HC | M | 42.85 | 10.35 | 8.70 | 0.64 | 6.14 | 5.98 | 4.79 | 0.22 | 3.02 | 2.86 | 2.73 |
| **POOL10** | HC | M | 55.07 | 8.61 | 5.44 | 0.51 | 4.47 | 4.49 | 3.77 | 0.07 | 0.86 | 0.86 | 0.99 |
| **POOL11** | HC | F | 41.66 | 7.54 | 5.22 | 0.39 | 4.25 | 4.19 | 2.73 | 0.06 | 0.78 | 0.74 | 0.78 |
| **POOL12** | HC | F | 40.37 | 10.86 | 9.09 | 0.52 | 6.74 | 6.61 | 4.65 | 0.17 | 2.52 | 2.46 | 2.41 |
| **ACIDOLA** | - | - | 2.46 | 96.67 | 96.66 | 13.89 | 92.47 | 95.03 | 93.83 | 13.89 | 92.37 | 94.84 | 93.83 |
| **BLACTIS** | - | - | 2.11 | 99.21 | 99.21 | 11.13 | 98.14 | 98.45 | 97.99 | 11.13 | 98.13 | 98.43 | 97.99 |
| **ARTIFICIAL** | - | - | 50.00 | 11.20 | 10.33 | 2.62 | 2.80 | 4.41 | 9.32 | 2.61 | 2.63 | 3.90 | 8.14 |

| taxon - genus | RR1 | RR2 | RR3 | RR4 | SP1 | SP2 | SP3 | SP4 | HC1 | HC2 | HC3 | HC4 | ACIDOLA | BLACTIS |
|---|---|---|---|---|---|---|---|---|---|---|---|---|---|---|
| 1912216 - Cutibacterium | 4 | 4.7 | 4.2 | 4.3 | 4.2 | 4.3 | 4.3 | 4.3 | 4.5 | 4 | 4.2 | 4.6 | 4 | 3.2 |
| 1386 - Bacillus | 3.3 | 4.2 | 3.9 | 3.7 | 3.9 | 4.7 | 4.1 | 4.2 | 4.2 | 3.6 | 3.6 | 4.3 | 4.3 | 3.2 |
| 1678 - Bifidobacterium | 2.8 | 4.1 | 4 | 2.5 | 3.1 | 5.6 | 4.1 | 3.9 | 4 | 2.5 | 2.3 | 3.1 | 3.2 | 7.5 |
| 1301 - Streptococcus | 3.3 | 4.8 | 3.3 | 3.4 | 3.6 | 5 | 3.7 | 4 | 4 | 3.5 | 3.6 | 4.7 | 3.1 | 1.5 |
| 1883 - Streptomyces | 3.1 | 3.9 | 3.5 | 3.5 | 3.6 | 3.9 | 3.7 | 3.6 | 3.9 | 3.3 | 3.3 | 3.9 | 3.2 | 2.2 |
| 1279 - Staphylococcus | 3.3 | 3.7 | 3.3 | 3.5 | 3.5 | 4.1 | 3.7 | 3.7 | 3.8 | 3.2 | 3.4 | 3.7 | 3.5 | 2.3 |
| 1578 - Lactobacillus | 2.8 | 3.6 | 2.9 | 3 | 3.5 | 4.1 | 3.3 | 3.5 | 3.5 | 3.2 | 4.2 | 3.4 | 7.5 | 3.3 |
| 32008 - Burkholderia | 2.9 | 3.4 | 3.4 | 3 | 3.4 | 3.8 | 3.6 | 3.6 | 3.8 | 3.2 | 3 | 3.6 | 2.6 | |
| 57495 - Dermacoccus | 2.5 | 3.8 | 2.6 | 3.5 | 4.1 | 3.8 | 3.1 | 3.2 | 3.6 | 3.4 | 3.5 | 3.4 | 2.4 | |
| 475 - Moraxella | 2.7 | 4 | 3 | 3.4 | 4.2 | 4.3 | 3.3 | 3.3 | 3.7 | 3.3 | 3.4 | 4.4 | 2.4 | 2.4 |
| 55193 - Malassezia | 3.2 | 3.8 | 3.5 | 3.4 | 3.2 | 3.4 | 3.5 | 3.2 | 3.7 | 3 | 3.1 | 3.5 | 2.8 | 2.3 |
| 374 - Bradyrhizobium | 3 | 3.4 | 3.2 | 2.8 | 3.2 | 3.4 | 3.7 | 3.3 | 3.5 | 3 | 3.1 | 3.6 | 2.7 | 1.6 |
| 5052 - Aspergillus | 2.7 | 3.4 | 2.9 | 3.2 | 3.4 | 3.3 | 3.3 | 3 | 3.7 | 2.8 | 2.9 | 3.4 | 2.9 | 1.5 |
| 570 - Klebsiella | 2.7 | 2.8 | 3.2 | 3.4 | 3 | 3.2 | 3.1 | 3.5 | 3.8 | 3.2 | 2.8 | 3.3 | 4.5 | 3.6 |
| 561 - Escherichia | 2.3 | 3.4 | 2.8 | 2.8 | 3.1 | 3.5 | 3 | 3.7 | 3.6 | 3 | 2.6 | 3.2 | 4.2 | 3.5 |
| 2701 - Gardnerella | 2.3 | 3.3 | 3 | 2.9 | 3 | 3.3 | 3.2 | 3 | 3.3 | 2.8 | 2.9 | 3.5 | 1.3 | 2.5 |
| 80865 - Delftia | 2.7 | 3.2 | 3 | 2.6 | 2.8 | 3.1 | 3.2 | 3.8 | 3.3 | 2.9 | 2.8 | 3.3 | 2.2 | |
| 590 - Salmonella | 2.1 | 3 | 2.6 | 2.5 | 2.8 | 3.1 | 2.9 | 4.1 | 3.3 | 2.8 | 2.3 | 3 | 4 | 3.2 |
| 44249 - Paenibacillus | 2.3 | 2.9 | 3.2 | 2.3 | 2.9 | 4.5 | 3.2 | 3.6 | 3.1 | 2.3 | 2.5 | 2.8 | 2 | |
| 2034 - Curtobacterium | 2.4 | 3 | 2.6 | 2.4 | 2.8 | 3.1 | 2.7 | 2.6 | 3.1 | 2.5 | 2.4 | 2.9 | 1.7 | |
| 68287 - Mesorhizobium | 1.8 | 3 | 2.5 | 2.4 | 2.7 | 2.8 | 2.7 | 2.5 | 2.7 | 2.3 | 2.7 | 2.7 | 2 | |
| 482 - Neisseria | 2.6 | 3.4 | 2.2 | 2.5 | 2.7 | 3 | 2.5 | 3 | 2.9 | 2.3 | 2.5 | 4.2 | 1.8 | |
| 53335 - Pantoea | 1.3 | 2.4 | 2.5 | 2.3 | 2.7 | 2.6 | 2.5 | 3 | 2.6 | 2.3 | 2.2 | 2.7 | 2.7 | 2.2 |
| 84567 - Pedobacter | 2.1 | 2.5 | 2.5 | 2.6 | 2.8 | 2.7 | 2.6 | 2.7 | 2.9 | 2.3 | 2.5 | 2.8 | 1.7 | |

Figure S2: Heatmap showing genera that did not pass the cut criteria of being 10 times more abundant in samples than in controls, with normalisation.

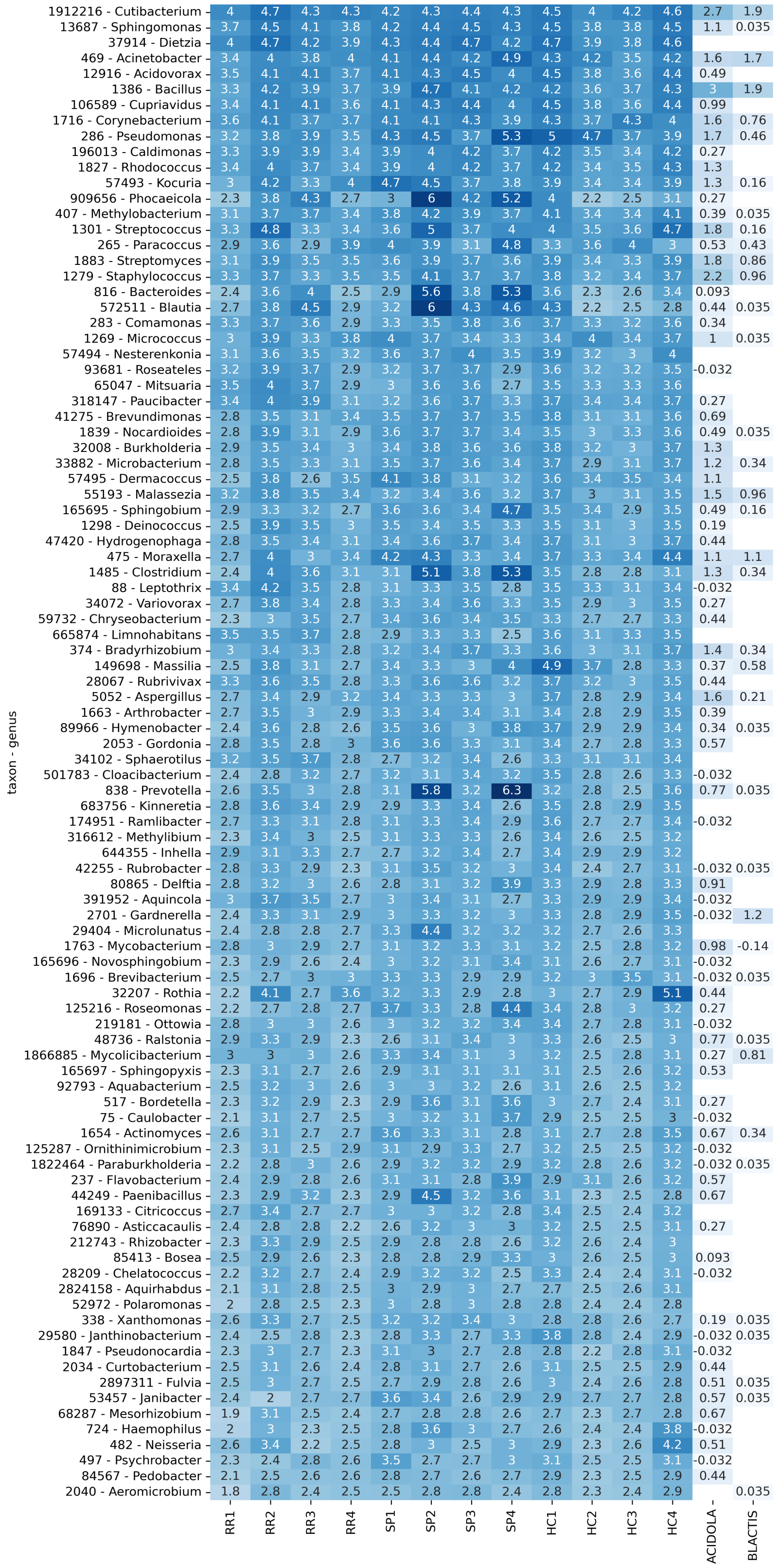


Figure S3: Heatmap showing genera that pass the cut criteria of being 10 times more abundant in samples than in controls, without normalisation.

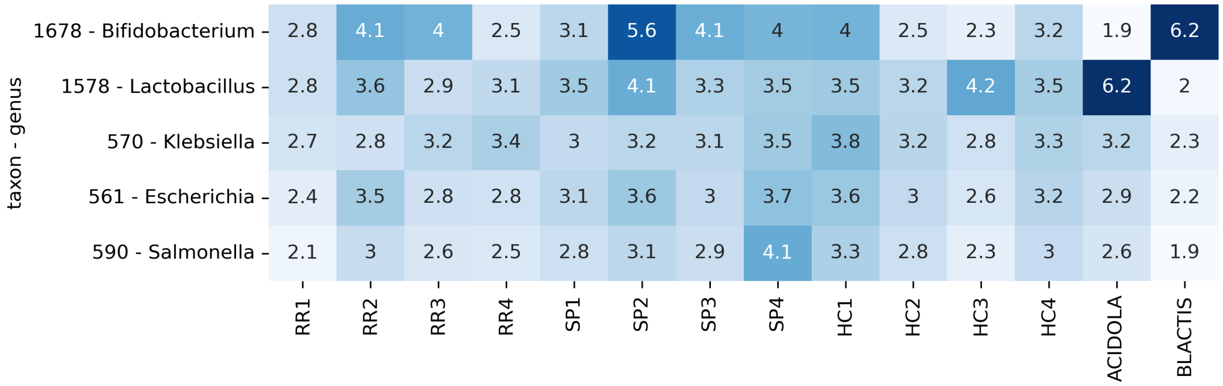


Figure S4: Heatmap showing genera that did not pass the cut criteria of being 10 times more abundant in samples than in controls, without normalisation.

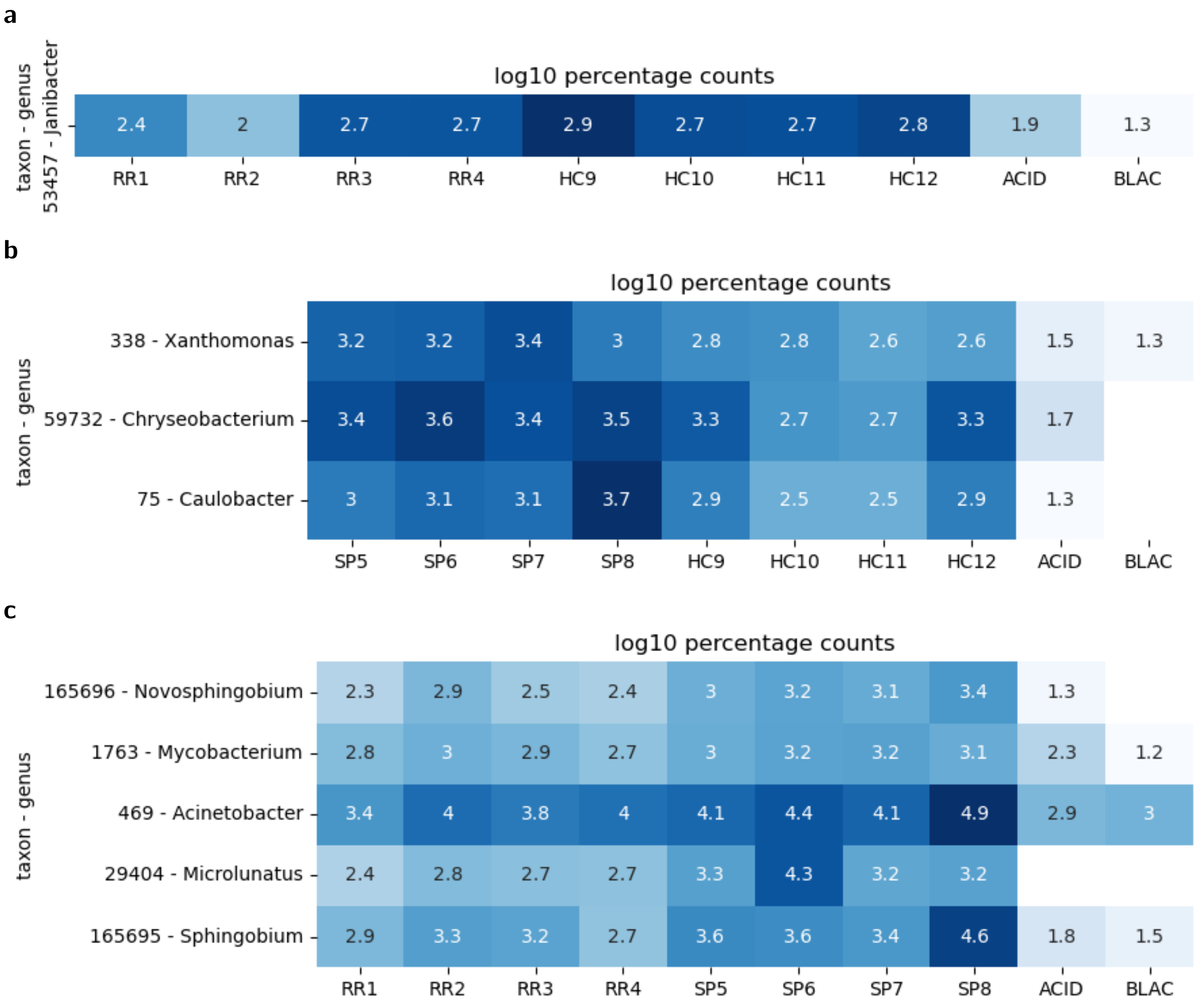


Figure S5: Heatmaps of counts assigned to differentially abundant genera, comparing (a) RR vs HC, (b) SP vs HC and (c) RR vs SP. Two-sided Mann-Whitney U test was applied ($p < 0.05$)
